# Recurrent neuronal loops between medial prefrontal cortex and ventral tegmental area display sex-specific spatial reorganization in response to stress

**DOI:** 10.64898/2026.03.23.713492

**Authors:** L. Pancotti, E. Dumas, A. Marroquin Rivera, C.D. Proulx, B. Labonté

**Affiliations:** CERVO Brain Research Center, Québec, QC, Canada; Department of Psychiatry and Neuroscience, Faculty of Medicine, Université Laval, Québec, QC, Canada; NeuroQuébec Thematic Research Center

**Author notes:** Corresponding author: Benoit Labonté.

**Keywords:** medial prefrontal cortex, ventral tegmental area, bidirectional connectivity, acute and chronic stress, sex-specificity, pathways-specific molecular profiling, functional clustering

## Abstract

The medial prefrontal cortex (mPFC) and ventral tegmental area (VTA) form a highly interconnected circuit involved in emotional regulation, stress reactivity, and cognitive processing. While prior research has established the anatomical and functional interactions between these regions, the precise organization and molecular identity of VTA neurons involved in unidirectional and bidirectional mPFC connectivity remains poorly defined, particularly under stress.

We combined dual anterograde and retrograde viral tracing in male and female mice to label VTA neurons according to their connectivity with the mPFC. This approach identified three distinct subpopulations including mPFC-projecting, mPFC-receiving, and bidirectionally-connected neurons which accounted for nearly half of the labelled VTA population. Each group displayed molecular heterogeneity, with most cells expressing dopaminergic (TH) and glutamatergic (VGLUT2) transcripts rather than single dopaminergic or GABAergic (GAD1) markers.

Acute and chronic stress exposure revealed sex- and circuit-specific patterns of c-Fos activation. In males, acute and chronic stress generated opposing rostrocaudally organized activation profiles, whereas females showed a more uniform increase in activity. Spatial clustering analyses further revealed that stress induces distinct hotspot organization within the VTA, with chronic stress promoting cohesive hotspot organization and consistent local enrichment of bidirectionally connected neurons despite a limited global activation.

Together, these findings uncover a molecularly diverse mPFC–VTA circuitry with bidirectional connectivity that undergoes sex-dependent spatial and functional rearrangement under stress, providing new insights on circuit-level mechanisms of stress-related disorders.

## INTRODUCTION

The medial prefrontal cortex (mPFC) is a central regulator of stress responses, emotional processing, and cognitive functions through its connections with limbic and midbrain structures [1–3]. Within this framework, the mPFC receives nearly half of its dopaminergic afferents from mesocorticolimbic neurons in the ventral tegmental area (VTA), supplemented by GABAergic and glutamatergic tegmental inputs [4,5]. These mesocortical afferents form a sparse but organized network across prelimbic and infralimbic regions [6], regulating higher-order functions including aversive processing, attentional control, and reinforcement learning [7–10].

Reciprocally, the mPFC sends dense glutamatergic projections to the VTA [11–13], arising from pyramidal neurons in cortical layers V and VI [14]. These cortical afferents converge onto a heterogeneous neuronal population in the VTA, including both dopaminergic and GABAergic neurons [11,15,16]. Functionally, VTA-projecting mPFC neurons are implicated in reward encoding, social hierarchy, and avoidance of aversive outcomes[7,17,18]. Together, this supports a bidirectional mPFC–VTA circuit in which ascending and descending pathways contribute to coordinated oscillatory activity associated with working memory, decision-making, and risk evaluation, as suggested by previous functional studies[17,19–21]. Given the specialization of mesocortical terminals for valence processing[6,22], identifying VTA neurons that both receive and provide mPFC input becomes critical. Defining whether these bidirectionally connected neurons constitute a distinct population is essential for uncovering the integrated logic of the mPFC–VTA circuit.

Stress induces major functional and morphological changes in both regions [23,24]. At the circuit level, chronic variable stress produces molecular and morphological alterations in mPFC neurons projecting to the VTA, accompanied by increased excitatory synaptic drive and a shift in synaptic E/I balance in both male and female mice [25]. In parallel, chronic social defeat stress has been shown to reduce dopaminergic axon density in the mPFC while increasing c-Fos tegmental activity in females [26], supporting the idea that stress alters both cortical output and mesocortical input. Consequently, bidirectionally connected mPFC–VTA neurons may represent a critical substrate through which stress reorganizes local circuit dynamics.

Here, we combined anterograde and retrograde trans-synaptic viral tracing to characterize VTA neuronal subpopulations based on their directional connectivity with mPFC. We identified three subpopulations, including a large subset of bidirectionally connected neurons. We then characterized their molecular signatures, assessed their recruitment following acute and chronic stress in males and females and evaluated their spatial reorganization throughout stress chronicity.

## METHODS

### Animals

All experiments were carried out on male and female C57BL6/N mice aged 8-12 weeks (Charles River). Mice were housed 2-4 per cage and kept on a 12/12 h light/dark cycle with food and water provided *ad libitum*. Experiments were conducted in accordance with the Canadian Guide for the Care and Use of Laboratory Animals and were approved by the Université Laval Animal Protection Committee.

### Stereotaxic surgery and viral strategy

Mice (80-90 days old) were anesthetized with isoflurane for bilateral stereotaxic injection of adeno-associated viruses (AAVs) in the mPFC and VTA. Injections were performed using a glass pipette mounted on a stereotactic frame, with volumes ranging from 150-200 nl per site. AAVs were infused at a rate of 5 nanoliters per second. Following each injection, the glass pipette remained at the injection site for 5 minutes to allow viral diffusion. For each experiment, mice were injected with a mix (1:1) of AAV1-hSyn-Cre-WPRE and AAVretro-EF1a-FlpO in the mPFC (1.8 AP; ±0.75 ML; -2.2 DV; 10°Angle) and a mix (1:1) of AAV5-hsyn-DIO-eGFP and AAV5-hsyn-fDIO-mCherry in the VTA (-3.3 AP; ±0.5 ML; -4.5 DV). AAVs used in this study, along with providers and titers, are listed in **Supp. Table 1**.

### Immunohistochemistry

Four weeks following viral injections, mice were deeply anesthetized with urethane (10mg/ml, intraperitoneal administration) and perfused by transcardial injection of 0.1M phosphate buffer (PB, p.H. 7.4) followed by 4% Paraformaldehyde (PFA) dissolved in PB. Brains were extracted and left overnight in 4% PFA before being transferred to a 10% sucrose solution in PB. Brains were sectioned at 50 μm with a LEICA VT1000S vibratome in the coronal plane and maintained at +4°C in PB.

IHC was used to identify the molecular identity of mPFC-VTA connected cells (Thyrosine Hydroxylase: TH+ and Dopamine Decarboxylase: DDC+) and to evaluate neuronal activity via c-Fos staining using the same protocol. A total of 11 male mice were used for TH and DDC experiments. More details on c-Fos experiments are provided in the neuronal activity section below. Free-floating tissue slices were washed with PB and incubated for 2 hours at room temperature in a blocking solution containing 10% Normal Donkey Serum (NDS), 3% Bovine Serum Albumine (BSA) and Triton 0.5%. Tissue slices were first incubated in a primary solution containing the primary antibody (anti-TH (1:500), anti-DDC (1:1000), anti-c-Fos (1:1000) diluted in PB), 1% NDS, 3% BSA and 0.5% Triton-X at +4°C overnight, before being incubated for 2 hours at room temperature in a solution containing the secondary antibodies. Slices were finally washed with PB and mounted on glass slides with Fluoromount medium (Sigma Aldrich, F4680). Primary antibodies for DDC (rabbit anti-DDC (1:1000), ab1569, Thermofisher) and c-Fos (rabbit anti-c-Fos (1:1000) ab214672, Abcam) were coupled with an Alexa Fluor 647 (anti-rabbit (1:1000), A315773, Thermofisher). TH (sheep anti-TH, (1:500), ab1541, Millipore) was incubated with Alexa Fluor 647(anti-sheep (1:1000), SAB4600178, Sigma-Aldrich).

### *In situ* hybridization

The intersectional viral approach described above was combined with RNAscope ISH to determine the molecular signature of the three different mPFC-VTA connected neuronal populations. A total of 9 male mice were used in these experiments. Four weeks following viral infections, mice were perfused with 4% Diethyl pyrocarbonate (DEPC)-PFA before brain extraction. Brains were kept in 4% DEPC-PFA overnight before being transferred to 30%-sucrose until sinkage. 30 μm free-floating VTA slices were kept in antifreeze solution at -20°C prior to ISH.

Free-floating sections were mounted on Superfrost Plus slides on the experimental day. The manufacturer’s pretreatment phase protocol (ACD – RNAscope Multiplex Fluorescent Reagent Kit v2) was adapted to account for slice thickness and optimize tissue permeabilization. Briefly, a washing phase was added on free floating sections using TRIS phosphate buffer containing 0.1% of Tween-20 (1X TBS-T 0.1%) prior to mounting the tissue slices on slides. An additional 1h tissue post-fixation period in fresh PFA (4%) was performed after the first 60°C incubation step. The ethanol gradient step was followed by a 15-minute incubation period at 60°C. Finally, the target retrieval period was performed for 5 minutes.

TH-mRNAs with either GAD-mRNAs or VGLUT2-mRNAs were co-labelled along the anteroposterior (AP) axis (from Bregma: rostral (-2.9), medial (-3.1), caudal (-3.3)) of the VTA. The TH RNAscope probe (Mm-Th-C3, 317621-C3, ACD) was coupled to the 480 fluorophore (Opal Polaris 480 Reagent Pack, FP1500001KT, Akoya). The GAD1 (Mm-GAD1, 400951, ACD) and VGLUT2 (Mm-Slc17a6, 319171, ACD) RNAscope probes were coupled to the 690 fluorophore (Opal 690 Reagent Pack, FP1497001KT, Akoya). Probes were provided by ACD (advanced cell diagnosis). RNAscope mRNA-staining steps were performed following the manufacturer’s protocol. A post-RNAscope IHC was performed to amplify the GFP and mCherry fluorescent signals induced by our viral approach. Primary antibodies (rabbit anti-GFP 1:1000, ab290, abcam, and rat anti-mCherry 1:1000, M11217, ThermoFischer Scientific) and secondary antibodies (Alexa goat anti-rabbit Alexa Fluor 488 1:1000, A11008, ThermoFischer Scientific, and Alexa 568 goat anti-rat Alexa Fluor 568 1:1000, A11077, ThermoFischer Scientific) were added to a blocking solution containing 5% of goat serum in DEPC-PBS-T 0.1%. Tissue slices were mounted on glass slides with mounting media and kept in the dark at 4°C until imaging. We extracted 200 neurons on average per mouse, from a total of 9 mice (TH-VGLUT experiment: N = 3; TH-GAD: N = 6).

### Image acquisition and cell count

Confocal images were acquired using a Nikon A1R HD microscope equipped with a 25× water-immersion objective. For each slice, a 3×2 tile scan of the VTA was taken in combination with Z-stack acquisition. Images were processed in FIJI (ImageJ), and three slices per animal, each corresponding to a different anterior-posterior (AP) position, were analyzed.

Signal colocalization of GFP^+^ and mCherry^+^ neurons was annotated using the Colocalization Object Counter plugin [27]. Colocalization was further confirmed by manual inspection of Z-stacks to ensure overlap occurred within the same focal plane. Proportions were calculated as the number of marker-defined cells over the total number of counted cells per slice and averaged across animals. To check if the observed proportions differed from the expected proportion assuming equal distribution of the three markers (1/3), we performed three independent chi-square tests for equality. For *in situ* hybridization analysis, colocalization of tyrosine hydroxylase (TH) with either VGLUT2 or GAD1 was assessed in GFP^+^, mCherry^+^, and dual-labeled (GFP^+^/mCherry^+^) neurons. For c-Fos quantification, confocal Z-stacks were processed in FIJI to generate maximum intensity projections. The VTA was manually masked, and c-Fos^+^ cells were identified using the 3D Objects Counter plugin with manually optimized intensity thresholds. This analysis yielded both total cell counts and XY coordinates, which were used for subsequent spatial analysis.

### Acute/Chronic stress paradigms

A single session of foot shock was used to induce acute stress in males and females as described before [28]. On experimental days, control mice were placed in the shocker cage for 60min without receiving any shock while a separate group of mice experienced a single 60min/100 random shocks session. Every mouse was sacrificed through a terminal urethane injection and perfused 60 minutes following the shocking session as described in the IHC section.

In parallel, a second group of male and female mice underwent chronic variable stress (CVS) as described before [25,29,30]. Briefly, CVS consists of three different stressors repeated for 21 days after which both males and females exhibit a range of behavioral alterations including anhedonia, behavioral despair, and anxiety. On the first day, mice are placed in a shocker where 100-foot shocks at 0.45 mA are randomly applied over one hour. On day two, mice are suspended by the tail for one hour. On the third day, mice are restrained in a 50mL Falcon tube for one hour. One day after the last stressor (day 22), mice received a final foot shock session (60min/100 random shocks) and were sacrificed 60 min later, as in the acute stress group. Control male and female mice were exposed to a shocker cage without shock 2 days prior to perfusion for novelty habituation.

In total, 32 mice (17 males and 15 females) were used in these experiments and divided in the following groups: control (Male, N=5; female, N=4), acute stress (Male, N=6; female, N=5) and chronic stress (Male, N=6; female, N=6). These experiments were performed on three cohorts for males (First cohort: 6; Second cohort: 5; Third cohort: 6) and two cohorts for females (First cohort: 9; Second cohort: 6).

### c-Fos absolute/cell-specific analysis

Neuronal activity in the VTA and its mPFC-connected neuronal populations was estimated following acute and chronic stress using c-Fos IHC, as described above. Variations in the absolute number of c-Fos^+^ neurons between stress conditions, across AP positions were assessed separately for each sex using Two-Way ANOVA with Sidak’s multiple comparison test. Significance was fixed at p<0.05. Analyses were performed with GraphPad Prism 10 (GraphPad Software, San Diego, CA).

For population-specific analysis, descriptive statistics were first performed for all variables according to their measurement scale. To account for variability related to batch membership while preserving meaningful between-group comparisons, counts were normalized to the batch A control group within each sex and marker population. The distribution of normalized c-Fos ratios was visually assessed and compared with theoretical simulations generated from different statistical distributions. Given the positive skew of the data, the measurement scale and the repeated measurements obtained from 3 AP positions per subject, Generalized Log-Linear Mixed Models were implemented. The final model, selected using likelihood ratio tests, included c-Fos ratio as the dependent variable, fixed effects for condition, sex, AP Position, marker population and interactions with condition, as well as random intercepts for AP position within subject. Deviation coding was applied to AP position and marker population because of their nominal nature, allowing estimation of simple effects when ratios were equal to the mean of these variables and their deviations around the grand mean. Because marker population accounted for the largest portion of variance in the model, additional models were fitted separately for each population while retaining all interactions as fixed effects. These statistical analyses were performed in the R environment using the lme4, lmerTest, MASS, DHARMa, gtsummary and ggplot2 packages.

### Clustering analysis

A multi-step spatial analysis was performed to compare the spatial distribution of c-Fos^+^ neurons across stress conditions using Local Indicators of Spatial Association (LISAs), a statistical method previously applied in neuroscience research [31,32]. Briefly, segmented neuron coordinates were pooled within each condition, normalized to a common [0,1] space, and aligned across animals. Kernel density estimation (KDE) was applied to generate population-level density maps and compute ACT-CTRL and CVS-CTRL difference heatmaps. Hotspot analysis was performed using Getis-Ord Gi* local spatial statistics (KNN = 10, 999 permutations) based on local density within a radius of 0.05. Significant hotspots (p<0.05) were subsequently refined using Local Moran’s I analysis to remove spatial outliers. Hotspot clusters were then identified using DBSCAN (ε = 0.07, min_samples = 10) and cluster properties including size, area and density were extracted. Group-level comparisons included centroid displacement (Euclidean distance between group centroids) and Jaccard Index. All results were mapped onto a standardized atlas to verify anatomical specificity. To quantify spatial overlap between hotspot regions, we computed a radius-based Jaccard index using one-to-one point matching within a spatial tolerance of 0.05. For each point in dataset A, the nearest unmatched point in dataset B within the given radius was counted as an intersection. The final Jaccard index was defined as the ratio of matched pairs to the union of all candidate points preventing overcounting through unique point assignment.

### Cluster Enrichment Analysis

Cell counts for each population were obtained from spatially defined activity clusters across anterior–posterior (AP) slices. Cell identities were derived from manually annotated fluorescently labeled neurons (e.g., GFP, mCherry, GFP/mCherry, and their respective c-Fos subtypes), using the same dataset used for c-Fos analyses. For each animal, the number of cells of each type was quantified within cluster and non-cluster regions. Counts were averaged by experimental condition (ACT, CVS) and sex (male, female). Enrichment ratios were calculated by comparing the proportion of each cell type within clusters relative to their proportion in the non-clustered portion of the same slice. The resulting ratios were then log₂-transformed and plotted as fold-change enrichment values. Positive values indicate selective enrichment within clusters, while negative values reflect depletion. To estimate the probability of observing equivalent or more extreme fold changes, reporter labels were shuffled and fold changes in proportions were recalculated to generate null distributions (10000 permutations). The p-value corresponds to the two-sided probability of occurrence of the observed values.

## RESULTS

### Combined anterograde and retrograde viral tracing revealed distinct connectivity patterns among VTA neurons linked to the mPFC

We first assessed mPFC-related connectivity patterns in VTA neurons using a dual intersectional viral strategy combining anterograde AAV1 and retrograde AAVs (**Figure 1A**). This approach was designed to define connectivity-related VTA populations while accounting for the known retrograde properties of AAV1 at titers used for anterograde tracing[33]. Specifically, we co-injected AAV1-hSyn-Cre (anterograde) and AAVrg hSyn-Ef1a-Flpo (1:1 ratio) into the mPFC. Simultaneously, a combination of conditional AAVs was injected in the VTA to drive the conditional expression of eGFP and mCherry in VTA neurons. Using this strategy, we identified three distinct populations linked to the mPFC: neurons receiving direct input from the mPFC were labeled with GFP, whereas mPFC-projecting VTA neurons expressed mCherry. Neurons co-expressing GFP and mCherry were classified as bidirectionallly connected, consistent with receiving inputs from and projecting back to the mPFC (**Figure 1A, B**).

**Figure 1.**
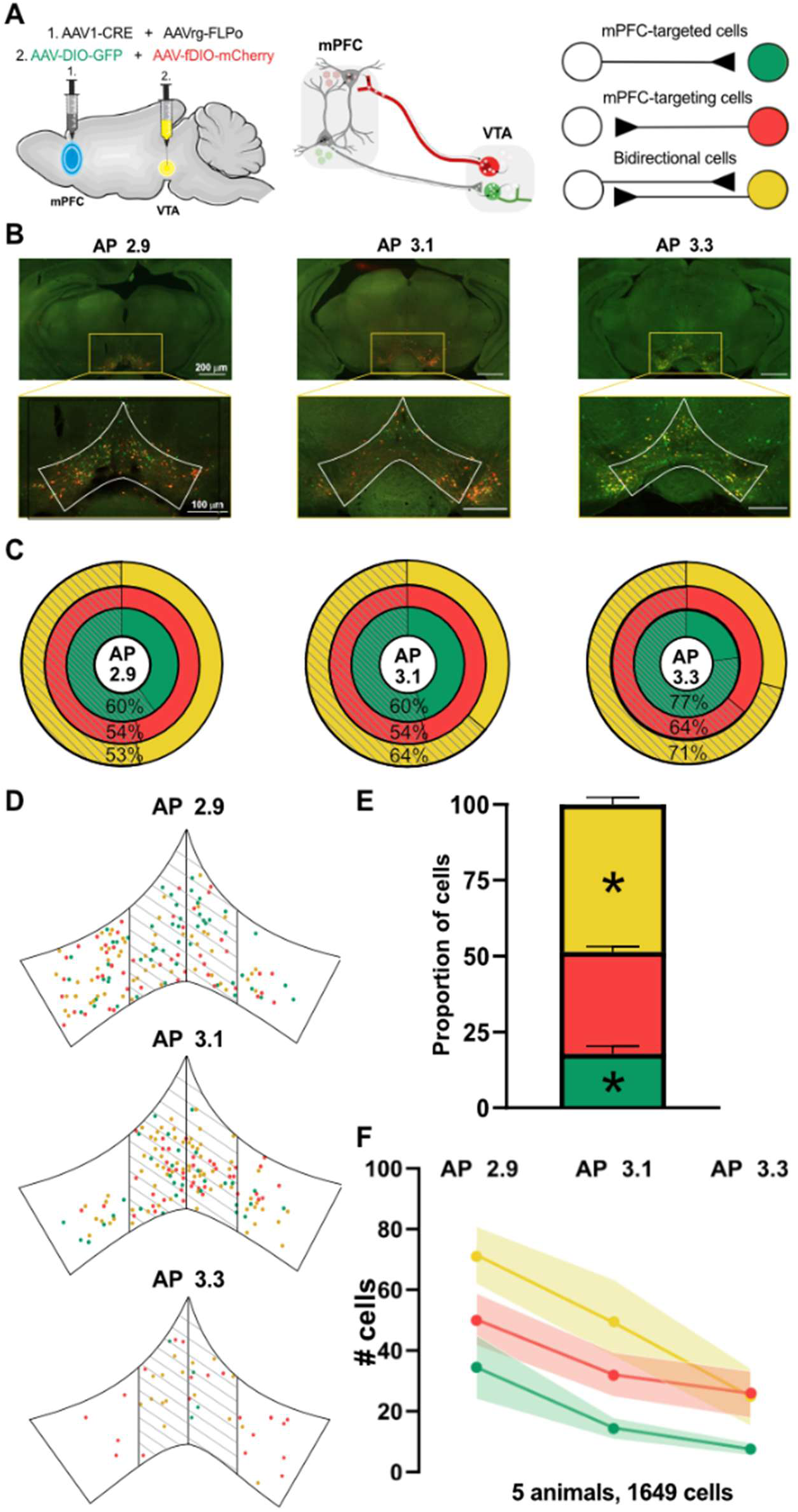
The mPFC and VTA exhibit robust bidirectional connectivity. **A** Schematic representation of the AAV intersectional strategy used to identify VTA neurons according to their connectivity with the mPFC. Three possible outcomes are depicted based on fluorescence expression. **B** Representative images of VTA sections across AP positions (Distance from Bregma: -2.9, -3.1, -3.3). Insets show representative VTA regions containing GFP, mCherry and GFP/mCherry cells. **C** Pie charts showing the medial versus lateral distribution of GFP, mCherry, and GFP/mCherry cells across the AP axis. **D** Representative distribution of AAV-tagged cells showing average distribution and lateralization across AP positions. **E** Proportion of labelled cells according to fluorescent identity. Data are represented as mean ± SEM. **F** Absolute number of labelled cells for each fluorescent population along the AP axis. Continuous lines indicate group mean and shaded areas represent SEM.

In total, we counted 1,649 fluorescently labeled cells from 5 mice. Notably, nearly half of these neurons (48.6%) co-expressed GFP and mCherry, indicating a large bidirectionally labeled population. Conversely, 33.6% of labeled neurons expressed only mCherry, and 17.8% expressed only GFP (**Figure 1E**). A χ² goodness-of-fit analysis showed that the proportion of co-expressing neurons was significantly higher than expected by chance (∼33.3%; χ²:131.23, p<0.0001), while GFP-only neurons were significantly less frequent than expected (χ²:145.99, p<0.0001). These findings suggest that a substantial proportion of VTA neurons receiving direct mPFC inputs also project back to the cortex, forming a bidirectional circuit.

We next evaluated the spatial distribution of these populations along the anteroposterior and mediolateral axes of the VTA. All three subpopulations were broadly distributed throughout the VTA (**Figure 1C, D, F**), although our analysis revealed a rostrocaudal gradient, with a higher density of labeled cells in the rostral VTA (**Figure 1F**). No discernible pattern was observed along the mediolateral axis overall. In rostral regions, neurons were evenly distributed between medial and lateral portions, comprising approximately 60% of GFP-labeled cells, 54% of mCherry-labeled cells, and 53% of co-labeled cells located medially. In contrast, in more caudal regions, all three subpopulations were preferentially localized closer to the midline, with 77% of GFP, 64% of mCherry, and 71% of co-labeled neurons clustered medially (**Figure 1C**).

Overall, our analysis revealed distinct VTA neuronal populations connected to the mPFC, including a substantial bidirectionally labeled subgroup. While labeled neurons were more prominent in the rostral VTA, infected neurons became progressively more concentrated around the midline in caudal sections.

### Molecular profiling of mPFC-VTA connected neurons

The VTA is roughly composed of 60% dopaminergic (DA), 35% GABAergic, and 5% glutamatergic neurons [34,35], with subpopulations co-expressing DA, GLUT or GABAergic markers have also been reported in various proportions along the antero-posterior and medio-lateral axes, and reports indicate that midbrain neurons can co-release dopamine and glutamate [36,37]. We next evaluated the molecular identity of these subpopulations to determine whether their molecular signatures reflected general organizational patterns in the VTA.

We first quantified the expression of dopaminergic markers tyrosine hydroxylase (TH) and dopa decarboxylase (DDC) in VTA neurons labelled via our trans-sectional viral approach (**Supp Figure 1).** Quantification showed that only 5% of each subpopulation was TH-positive and 14% was positive for DDC (**Supp Fig. 1A-B**). Given previous reports characterizing mPFC-projecting cells in the VTA[11], and given that AAV infection has been shown to downregulate dopaminergic markers’ expression in the VTA [38], we concluded that this approach underestimated dopaminergic proportions.

Consequently, we used RNAScope ISH to quantify the expression of TH, VGLUT2, and GAD1 mRNA in GFP^+^, mCherry^+^, and GFP^+^/mCherry^+^ neurons, allowing to detect AAV fluorescence and mRNA expression in the same tissue (**Figure 2A-B**). Our analysis revealed a distinct molecular gradient along the anteroposterior (AP) axis of the VTA, with varying proportions of DA (TH^+^), GLUT (VGLUT2^+^), and GABAergic (GAD1^+^) neuronal populations (**Figure 2B-E, Supp Table 2**). The rostral VTA contained a higher proportion of glutamatergic neurons (TH^-^/VGLUT2^+^), while the medial VTA was dominated by TH^+^/VGLUT2^+^ co-expressing cells (**Figure 2B**). In contrast, the caudal VTA had a predominance of purely dopaminergic (**Figure 2B**) and GABAergic neurons (**Figure 2D, E**), supposing functional specialization within mPFC-connected populations.

**Figure 2.**
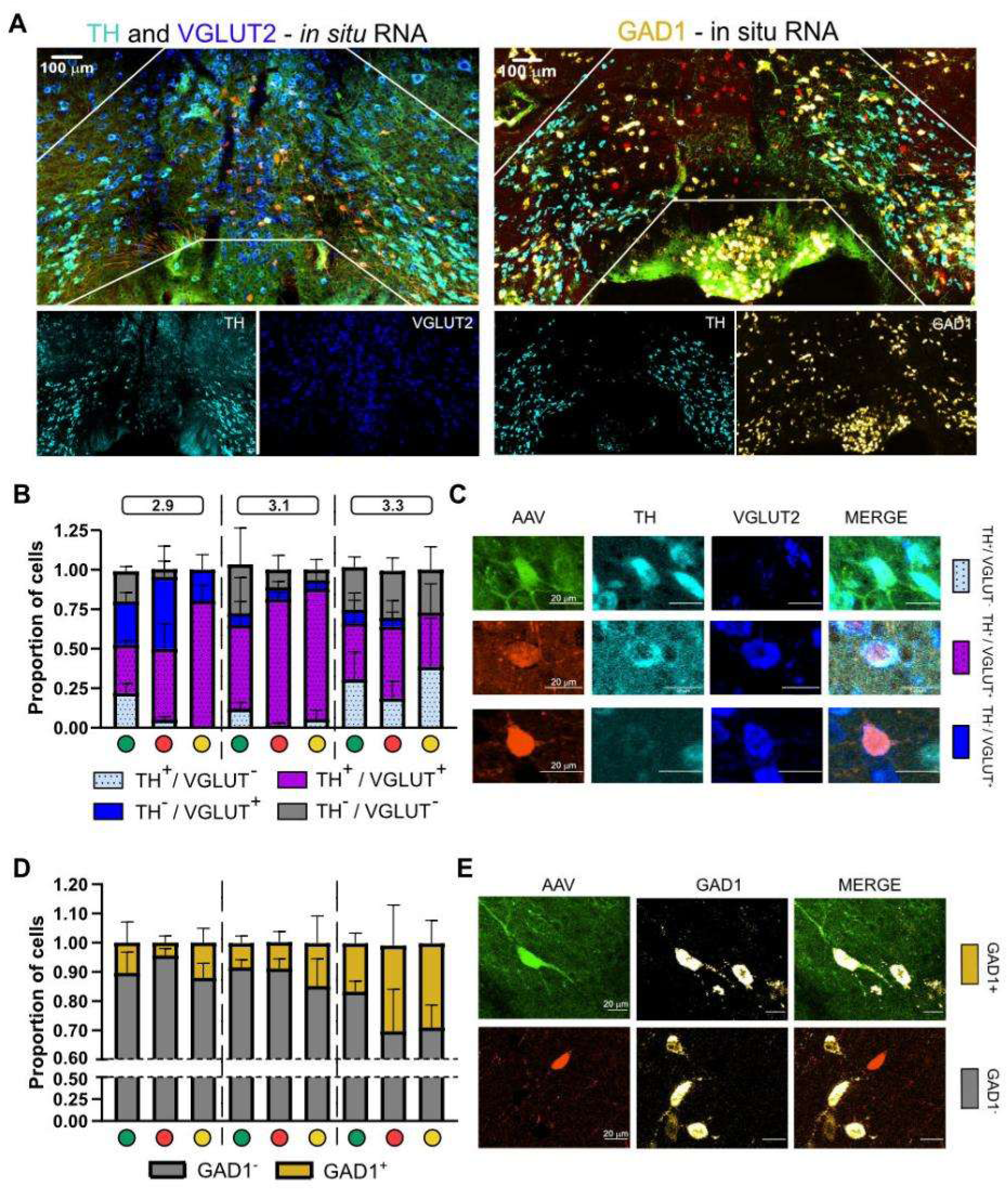
Molecular characterization of mPFC-VTA connected neuronal populations. **A** Representative images RNAscope ISH coupled with IHC staining for AAV fluorescent signal amplification. (**Left**) TH (cyan) and VGLUT2 (blue) labelling in VTA. (**Right**) TH (cyan) and GAD1 (gold) labelling in VTA. **B** Bar plot showing proportions of cells according to TH/VGLUT2 expression across fluorescent neuronal populations. Coloured dots below the X axis indicate the fluorescent populations. Boxes above indicate AP localization. Light blue: TH^+^/VGLUT2^-^; Purple: TH^+^/VGLUT2^+^; Blue: TH^-^/VGLUT2^+^; Gray: TH^-^/VGLUT2^-^. **C** Representative images and insets for each fluorescent population shown in panel B. **D**. Bar plot showing proportions of cells according to GAD1 expression across fluorescent neuronal populations. Gold: GAD1+. Gray: GAD1-. **E** Representative image and insets for fluorescent populations shown in panel D. Data are represented as mean ± SEM.

The GFP-only population displayed regional variations along the AP axis (**Figure 2B-D, Supp Table 2**). These cells were heterogeneously distributed in the rostral VTA, with a mix of VGLUT2^+^and TH^+^/VGLUT2^+^ co-expressing neurons. In the medial VTA, most GFP-positive cells co-expressed TH and VGLUT2, with minimal GAD1 expression (∼8%). In the caudal VTA, GFP-positive cells were evenly distributed between purely dopaminergic (TH-only) and TH^+^/VGLUT2^+^ co-expressing neurons, while the proportion of GAD1^+^ cells increased to approximately 16%.

The mCherry-only population also exhibited AP axis shifts (**Figure 2B, Supp Table 2**). In the rostral VTA, these cells were predominantly VGLUT2-only, indicating a strong glutamatergic phenotype. In the medial VTA, they displayed a higher degree of TH^+^/VGLUT2^+^ co-expression, with very few purely dopaminergic neurons. In the caudal VTA, about 50% of the mCherry-positive cells co-expressed TH and VGLUT2, while approximately 30% expressed GAD1, highlighting a mixed DA and GABAergic profile.

Finally, the GFP/mCherry population mirrored the mCherry-only population in the anterior and medial VTA, with roughly 80% of cells co-expressing both TH and VGLUT2, and almost no TH-only cells (**Figure 2B, Supp Table 2**). In the caudal VTA, however, this population showed a stronger GABAergic component, with nearly 30% of cells being GAD1-positive.

Together, our findings support substantial molecular diversity across VTA neuronal populations, with a predominance of VGLUT2/TH co-expressing neurons in rostral and medial sections and greater GABAergic presence caudally, suggesting that different functional subpopulations may contribute to mPFC-VTA connectivity and signaling.

### Stress-Induced Activation of mPFC-Connected VTA Neurons

We next assessed stress-induced recruitment of these populations by quantifying c-Fos expression following acute and chronic stress in both sexes (**Figure 3A**).

**Figure 3:**
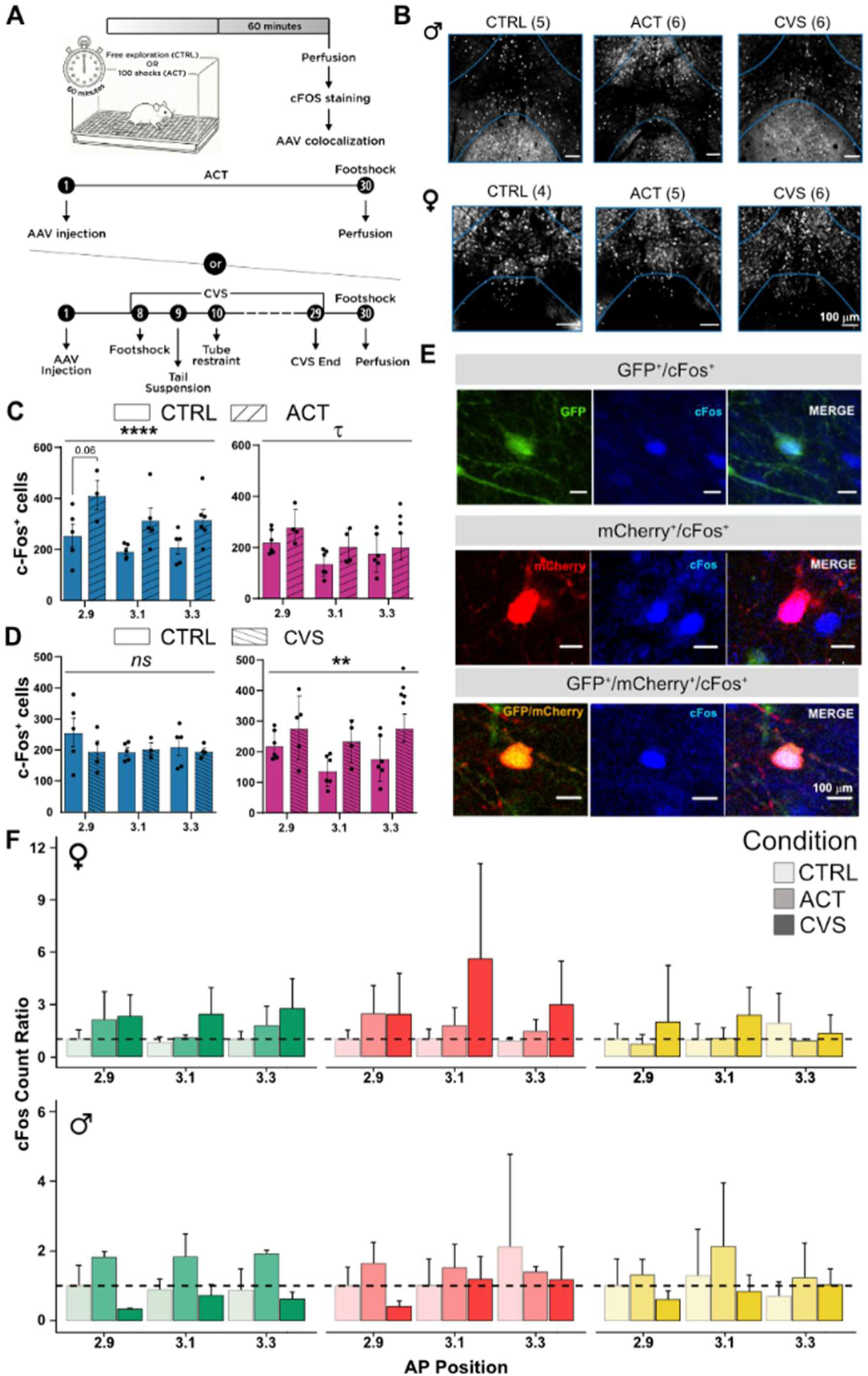
Acute and chronic stress differentially activates mPFC-VTA connected neuronal populations. **A** Experimental design of the c-Fos experiment, showing acute (ACT) and chronic variable stress (CVS) timelines. **B** Representative VTA sections across groups and sexes showing c-Fos staining (gray) in control, acute and chronic stress conditions. **C** Bar plots showing absolute number of c-Fos cells in control and acute conditions along the AP axis. Blue bars: males; pink bars: females. **D** Same analysis as in panel C for CVS condition. **E** Representative images of mPFC-connected fluorescently labelled neurons expressing c-Fos. **F** Normalized c-Fos ratios for each marker population according to condition, sex, and AP position. Ratios were normalized within each marker and sex to the mean control value at AP -2.9. The dashed line indicates the normalized control mean (1). Data are shown as mean ± SD. **** p<0.0001; ** p<0.01, τ p<0.1.

We first examined the effect of acute stress on c-Fos expression throughout the VTA (**Figure 3B, C**). Acute stress significantly increased c-Fos expression in males (F_(1,22)_=14.63, p<0.001), with a trend toward increased activation specifically in the rostral VTA (Post-hoc comparison, p = 0.06). In females, acute stress produced a trend toward a significant main effect (F_(1,10)_=4.21, p<0.07).

Notably, female c-Fos expression varied significantly along the AP axis (F_(2,14)_=4.66, p<0.03), with higher densities in rostral versus caudal regions.

We then tested whether CVS induces similar effects (**Figure 3B, D**). In males, CVS did not significantly alter c-Fos expression across the VTA. Conversely, females showed a significant increase in c-Fos expression after chronic stress (F_(1,10)_=12.04, p<0.006), with no significant main effect of AP position. This suggests distinct sex-specific activation patterns, with males and females responding more strongly to acute and chronic stress, respectively.

We next quantified c-Fos expression in GFP^+^, mCherry^+^, and GFP^+^/mCherry^+^ neuronal populations in the VTA (**Figures 3E-F**). Mixed-effect modeling (**Supp Table 3**) revealed that CVS nearly doubled c-Fos activation ratios across populations (Exp(β)=1.99, CI95%: 1.31 – 3.02, p< 0.005). While males showed higher baseline activation levels (Exp(β)=1.55, CI95%: 1.02 – 2.36, p<0.05), their response to CVS was attenuated, indicated by a significant stress-by-sex interaction (Exp(β)=0.39, CI95%: 0.22 – 0.72, p<0.005). Our model also revealed robust differences between neuronal populations (**Supp Table 3**).

Given the uniqueness of these cell types, we examined associations separately within each neuronal population (**Supp Table 4-6**). Within the GFP^+^ and mCherry^+^ populations, males displayed a bidirectional response: acute stress increased c-Fos expression, whereas CVS significantly reduced it. In females, both GFP⁺ and mCherry⁺ populations showed significantly increased c-Fos expression following acute and chronic stress. Notably, in both sexes, no significant changes were found within the bidirectional GFP+/mCherry+ neurons.

Overall, single-labeled populations exhibited the strongest stress sensitivity across sexes, while bidirectionally connected neurons remained stable across conditions.

### Global reorganization of mPFC-VTA connected neurons recruitment during acute and chronic stress

We next examined whether stress altered the spatial organization of activated neurons by comparing the spatial distribution of c-Fos expressing cells during acute stress and CVS (**Supp Figure 2**). After normalizing x and y coordinates into a common [0,1] space, we used kernel density estimation to generate population-level density maps (**Supp Figure 3**). Getis-Ord GI* identified statistically enriched hotspots, and DBSCAN clustering defined hotspot properties across VTA sections and experimental conditions (**Figure 4A, B**).

**Figure 4.**
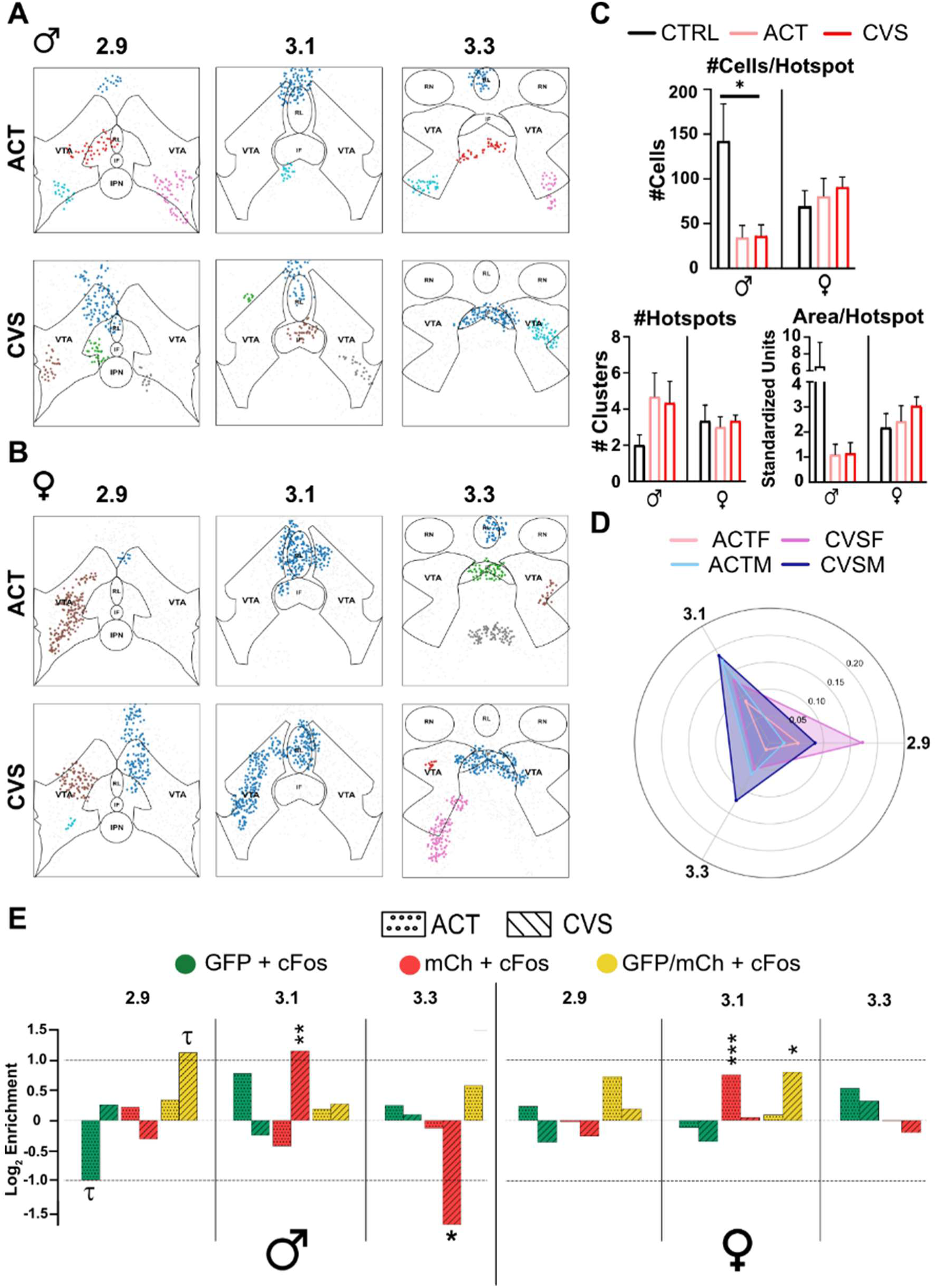
Spatial clustering of VTA c-Fos+ hotspots across stress conditions and sexes. **A–B** DBSCAN-based maps showing spatial distribution of c-Fos+ hotspots in (**A**) males under acute stress (ACT) and chronic variable stress (CVS), and (**B**) females under ACT and CVS conditions. Colors are arbitrarily assigned to c-Fos+ neurons belonging to distinct clusters. Hotspots are overlaid onto anatomical maps including the ventral tegmental area (VTA), interfascicular nucleus (IF), and Rostral Linear Nucleus (RL). **C** Quantification of hotspot properties across control, ACT, and CVS conditions in males and females, including the number of cells per hotspot, total number of hotspots, and total area covered by hotspots. **D** Radar plots showing the Jaccard index of spatial overlap between hotspots across experimental groups along the AP axis. **E** Log₂ fold change from enrichment analysis showing the proportion of each fluorescent cell population within clusters relative to the remaining portion of the slice. Positive values indicate enrichment and negative values indicate depletion. Dotted bars: ACT; hatched bars: CVS. Color codes indicate cell identity as shown in the legend. Categories with no available values are omitted. t p < 0.1; * p < 0.05; ** p < 0.01; *** p < 0.005.

In males, the number of c-Fos hotspots (2 to 5) did not differ significantly between groups (**Figure 4C**). However, the number of cells per hotspot was significantly reduced following acute and chronic stress (F_(5,12)_=3.25, p<0.05; **Figure C**), while hotspot area also showed a non-significant reduction under both conditions **(Figure 4C**). Acute stress generated fragmented clusters while CVS produced larger, more cohesive hotspots of c-Fos-expressing neurons, especially in the caudal VTA. For instance, male CVS produced a large midline cluster at -3.3 AP overlapping the interfascicular nucleus (IF), whereas acute stress clusters were mostly peripheral (**Figure 4A, B**). In females, hotspot analysis identified 3–4 clusters per group with no significant differences in frequency, cell density, or total area (**Fig. 4C**). Nevertheless, while acute stress hotspots were small and centered at -3.1 AP, CVS generated cohesive clusters throughout the AP axis (**Figure 4A, B**).

Finally, Hotspot overlap analysis revealed low Jaccard index (J) values across all groups, indicating that stress-induced distributions were distinct from baseline (**Figure 4D**). In males, acute stress produced minimal overlap rostrally (J = 0.03) but higher medial overlap (J = 0.18). Following CVS, overlap increased slightly in rostral and caudal sections (J = 0.08 and 0.12, respectively), whereas medial overlap remained stable (J = 0.19). In females, acute stress induced low overlap across all AP levels (J = 0.05, 0.09, and 0.01), while CVS increased overlap throughout the VTA, particularly rostrally (J = 0.17). Overall, males showed greater hotspot preservation in medial and caudal regions, while females exhibited stronger rostral overlap following chronic stress (**Figure 4D**). Together, these observations indicate that despite modest regional differences, hotspot overlap with control conditions remained globally low across all groups, supporting sex-and region-dependent changes in hotspot organization within the VTA.

Finally, we overlaid c-Fos hotspot maps with the topological distribution of AAV-tagged subpopulations. All three groups were detected within stress-associated clusters with differential rostrocaudal distribution under both stress conditions (**Supp Figure 4**). We assessed whether c-Fos^+^/labeled neurons were differentially represented within clusters relative to their overall abundance. In males, double-labeled/c-Fos^+^ neurons were consistently enriched within clusters under both stress conditions (**Figure 4E**, **Supp Figure 4**), with rostral CVS enrichment trending toward significance (p=0.08). GFP^+^/c-Fos^+^ neurons were generally enriched, except for a reduction observed in the rostral acute stress section (p=0.07, **Figure 4E**). In contrast, mCherry^+^/c-Fos^+^ neurons displayed region-dependent changes, with significant enrichment in medial CVS clusters (p<0.01) and significant depletion in caudal CVS clusters (p<0.05; **Figure 4E**, **Supp Table 4**).

In females, GFP^+^/c-Fos^+^ cells largely followed global c-Fos distributions, with only modest reductions at -3.1 under both stress conditions. mCherry^+^/c-Fos^+^ neurons similarly reflected global distributions, except for a marked enrichment at -3.1 following acute stress (p<0.01; **Figure 4E**, **Supp Table 4**). As in males, double-labeled/c-Fos^+^ neurons were consistently overrepresented within clusters when present, reaching significant enrichment under CVS at -3.1 (p<0.05; **Figure 4E**, **Supp Table 4**).

Together, GFP^+^/mCherry^+^ neurons were most consistently enriched within stress-associated hotspots, while GFP^+^ and mCherry^+^ populations displayed sex- and region-dependent local differences.

## DISCUSSION

The VTA and mPFC are highly interconnected brain regions whose bidirectional communication is critical for regulating complex cognitive and affective behaviors. Here, we further characterized the organization of mPFC-VTA connectivity and evaluated how distinct neuronal populations are affected by acute and chronic stress in males and female mice. Our findings support the existence of three major subpopulations of neurons including mPFC neurons projecting to the VTA, VTA neurons projecting to the mPFC, and a discrete subset of neurons exhibiting bidirectional connectivity. These populations exhibit distinct molecular signatures, with a predominance of neurons co-expressing dopaminergic and glutamatergic markers. Furthermore, they show unique spatial activation patterns in response to acute and chronic stress, suggesting functional heterogeneity within mPFC-VTA circuits.

Previous studies have characterized unidirectional connectivity between the mPFC and VTA, showing dense glutamatergic projections contacting the tegmentum from mPFC, and dopaminergic projections from the VTA innervating distinct layers of the mPFC [12,13]. Consistent with these findings, we identified both mPFC neurons projecting to the VTA and VTA neurons projecting to the mPFC. However, by combining anterograde and retrograde AAV tracing approaches, we also identified a population of VTA neurons that both receive input from and project to the mPFC. Although reciprocal VTA-mPFC loops have been previously suggested [11], our data expand this characterization by showing that nearly half of the VTA dopaminergic neurons receiving mPFC inputs also project to the cortex. These bidirectionally connected neurons follow a clear anteroposterior gradient within the VTA, with the highest density in rostral regions, consistent with previous anatomical descriptions [16,39], but without mediolateral organization. Bidirectional connectivity has been implicated in adaptive cognitive and affective regulation, including prediction error processing and flexible responses to changing contingencies [40,41], and may be essential for synchronizing mPFC-VTA activity during reward and aversion processing under stress [15,17,19,21,42].

The VTA is a neurochemically heterogeneous structure composed of dopaminergic, GABAergic, glutamatergic, and co-transmitting neurons [43,44]. Previous studies have shown that mPFC projections target dopaminergic and GABAergic neurons in the VTA [13]. Here, we extend these observations by characterizing the molecular identity of VTA neurons receiving mPFC input, projecting to the mPFC, or participating in bidirectional communication. Consistent with prior findings [36,37], we observed widespread TH/VGLUT2 co-expression across the VTA. This molecular profile was especially prevalent among bidirectionally connected neurons and VTA neurons projecting to the mPFC, with a strong medial distribution. In contrast, TH-only neurons were rare and confined primarily to caudal regions of the VTA. Consistent with previous reports [37,45], VGLUT2-only neurons exhibited a rostral bias, constituting 25–30% of VTA neurons receiving inputs from, and projecting to the mPFC. Finally, 20–30% of VTA neurons expressed GAD1 across the subpopulations, in line with recent transcriptomic studies identifying GABAergic, dopaminergic, and co-expressing subpopulations among PFC-connected neurons [16]. Although our marker set was limited, our findings support substantial neurochemical diversity among VTA neurons interacting with the mPFC, likely contributing to their functional specialization. Both the mPFC and VTA are strongly implicated in stress-related behaviors [23–26,46,47]. Stress-induced alterations in VTA dopaminergic neuron activity have been implicated in the encoding of aversive experiences [48] and susceptibility to depressive-like states [49,50], while chronic stress disrupts mPFC structural and functional organization [25,51], contributing to cognitive and emotional impairments. Here, we show that VTA neurons connected to the mPFC exhibit distinct recruitment patterns following acute and chronic stress, with marked sex-specific differences. In females, both receiving and projecting populations in the VTA were robustly activated by acute stress and remained responsive under chronic stress conditions. In contrast, bidirectionally connected neurons appeared largely unresponsive. In males, however, bidirectionally-connected neurons exhibited a biphasic pattern, being activated by acute but not by chronic stress, suggesting sex-specific adaptations of mPFC–VTA communication under prolonged stress exposure. Additionally, we observed consistent activation of VTA neurons projecting to the mPFC in response to acute stress in males, in line with previous studies demonstrating enhanced dopaminergic transmission to the mPFC under aversive and stressful condition [10,52]. While less is known about VTA neurons receiving mPFC inputs, our findings suggest they are strongly recruited during acute stress and disengaged following chronic stress exposure.

Contrary to expectations, overall c-Fos levels within the VTA did not markedly increase across stress conditions. A spatial hotspot analysis revealed distinct clusters of activity across groups, with a consistent medial bias in males and females. This organization is consistent with previous reports [53,54], showing that acute aversive stimuli inhibit lateral VTA dopamine neurons while preferentially activating the medial DA populations and now extends these observations by demonstrating similar organization in females. Importantly, acute and chronic stress generated distinct hotspot configurations, suggesting that stress duration may alter the spatial organization of activated neurons within the VTA. These clusters converged onto defined VTA subregions with known molecular and connectivity heterogeneity [55,56], suggesting that acute and chronic stress may selectively engage distinct components of mPFC-related VTA circuitry, the evolution of which may account for the transition from acute to chronic stress responses and sex-specific differences in stress susceptibility [57,58].

A particularly striking observation emerged from bidirectionally-connected neurons which showed limited recruitment in the global c-Fos analyses but were consistently enriched within stress-induced hotspots under both acute and chronic stress in males and females. This suggests that neuronal populations with minimal overall activation can exhibit focal activation, forming highly specialized functional clusters in response to stress, as previously suggested [59–61]. Overall, these findings suggest that stress does not simply alter global VTA neuronal activity but rather reorganizes how specific mPFC-connected neuronal subpopulations are engaged during stress.

Several methodological limitations should be acknowledged. First, AAV1 vectors used for anterograde tracing are known to exhibit limited retrograde properties [33,62]. Although prior work suggests that retrograde labeling remains minimal under similar experimental conditions [62], it remains possible that part of the bidirectional labelling observed here may be partially influenced by this characteristic. Future studies with more selective viral tools will be necessary to validate these connectivity patterns. Second, while c-Fos is a widely used proxy for neuronal activity, it remains an indirect marker, subject to variability in staining efficiency and cell type–specific expression dynamics[63,64]. While our findings are consistent with previous reports demonstrating stress-induced activation of mPFC and VTA circuits [25,48,65–67], functional validation using electrophysiological or imaging approaches will be required to confirm the behavioral relevance of these observations.

To conclude, our study reveals a complex and heterogeneous organization of mPFC–VTA connectivity, involving neuronal populations with distinct molecular signatures and connectivity profiles. We identified a subset of VTA neurons that both receive input from and project to the mPFC and show that these bidirectionally connected neurons exhibit spatially and sexually distinct activation patterns in response to acute and chronic stress. Together, these findings suggest that stress does not simply alter global VTA activity but selectively engages mPFC-connected neurons according to their molecular identity, anatomical location, and connectivity profile. Refined viral strategies combined with circuit-level functional analyses will be essential for dissecting the temporal dynamics and behavioral relevance of this bidirectional pathway.

## ACKNOWLEDGMENTS

The authors thank Patrick Desrosiers for his valuable input on the computational clustering analysis, Maryse Pinel, Chenqi Zhao, Modesto Peralta III, and Quentin Leboulleux for their essential technical assistance. LP, ED and AMR are supported by scholarships from the *Fonds de Recherche Quebec – Volet Santé*. BL holds a *Sentinelle Nord* Research Chair, is supported by the Canadian Institutes of Health Research (CIHR) (Grant No. PJT-451728 and PJT-451858) and the Natural Sciences and Engineering Research Council of Canada (NSERC) (Grant No. RGPIN-2019-06496) and receives Fonds de Recherche en Santé du Québec (FRQS) Senior salary support. CDP is supported by CIHR grant PJT-169117 and NSERC grant RGPIN-2017-06131.

## AUTHORS CONTRIBUTION

L.P. contributed to conceptualization, data curation, formal analysis, investigation, visualization, and manuscript writing and review. E.D. contributed to investigation, methodology, validation, and methods writing. A.M.R. performed formal statistical analysis, visualization, and manuscript writing. C.P. contributed to conceptualization, supervision, and manuscript review. B.L. contributed to conceptualization, funding acquisition, supervision, and manuscript writing and editing.

## Declaration of generative AI and AI-assisted technologies in the manuscript preparation process

During the preparation of this work the author(s) used Claude Sonnet 4.6 in order to proofread the manuscript text. After using this tool/service, the author(s) reviewed and edited the content as needed and take(s) full responsibility for the content of the published article.

---

**Supplementary Table 1.** Adeno-associated viral vectors.

| Serotype | Promoter | Construct | Provider | Lot | GC/ml |
| --- | --- | --- | --- | --- | --- |
| AAV1 | hSyn | CRE | Addgene | 885 | 2.4<br>E+13 |
| AAV2Rg | EF1a | FLP | Addgene | V56725 | 1.02<br>E+13 |
| AAV5 | hSyn | DIO-eGFP | Addgene | V54234 | 1.4<br>E+13 |
| AAV5 | EF1a | fDIO-mCherry | Addgene | V66588 | 1.1<br>E+13 |

**Supplementary Table 2.** Molecular characterization of neuronal populations expressing GFP, mCherry, or GFP/mCherry across the anteroposterior axis.

| **AP 2.9** | GFP | mCherry | GFP/mCherry |
| --- | --- | --- | --- |
| TH<sup>+</sup>/VGLUT2<sup>-</sup> | 0.22 ± 0.06 | 0.05 ± 0.02 | 0 |
| TH<sup>+</sup>/VGLUT2<sup>+</sup> | 0.30 ± 0.02 | 0.45 ± 0.16 | 0.80 ± 0.09 |
| TH<sup>-</sup>/VGLUT2<sup>+</sup> | 0.27 ± 0.05 | 0.45 ± 0.19 | 0.19 ± 0.09 |
| TH<sup>-</sup>/VGLUT2<sup>-</sup> | 0.09 ± 0.03 | 0.05 ± 0.05 | 0 |
| GAD1<sup>-</sup> | 0.90 ± 0.07 | 0.96 ± 0.02 | 0.88 ± 0.05 |
| GAD1<sup>+</sup> | 0.10 ± 0.07 | 0.04 ± 0.05 | 0.12 ± 0.05 |
| **AP 3.1** | GFP | mCherry | GFP/mCherry |
| TH<sup>+</sup>/VGLUT2<sup>-</sup> | 0.12 ± 0.04 | 0.01 ± 0.01 | 0.05 ± 0.05 |
| TH<sup>+</sup>/VGLUT2<sup>+</sup> | 0.53 ± 0.30 | 0.80 ± 0.04 | 0.82 ± 0.04 |
| TH<sup>-</sup>/VGLUT2<sup>+</sup> | 0.07 ± 0.07 | 0.07 ± 0.03 | 0.05 ± 0.05 |
| TH<sup>-</sup>/VGLUT2<sup>-</sup> | 0.31 ± 0.23 | 0.11 ± 0.09 | 0.06 ± 0.06 |
| GAD1<sup>-</sup> | 0.92 ± 0.02 | 0.91 ± 0.03 | 0.85 ± 0.09 |
| GAD1<sup>+</sup> | 0.08 ± 0.02 | 0.09 ± 0.15 | 0.15 ± 0.09 |
| **AP 3.3** | GFP | mCherry | GFP/mCherry |
| TH<sup>+</sup>/VGLUT2<sup>-</sup> | 0.31 ± 0.17 | 0.19 ± 0.11 | 0.40 ± 0.31 |
| TH<sup>+</sup>/VGLUT2<sup>+</sup> | 0.36 ± 0.19 | 0.45 ± 0.16 | 0.34 ± 0.18 |
| TH<sup>-</sup>/VGLUT2<sup>+</sup> | 0.08 ± 0.06 | 0.06 ± 0.03 | 0 |
| TH<sup>-</sup>/VGLUT2<sup>-</sup> | 0.27 ± 0.06 | 0.30 ± 0.08 | 0.27 ± 0.15 |
| GAD1<sup>-</sup> | 0.83 ± 0.04 | 0.70 ± 0.14 | 0.71 ± 0.07 |
| GAD1<sup>+</sup> | 0.16 ± 0.03 | 0.29 ± 0.14 | 0.29 ± 0.08 |

**Supplementary Table 3.** Linear-log model statistics.

| *Predictors* | **Ratios** |  |  |
| --- | --- | --- | --- |
|  | *Estimates* | *CI* | *p* |
| (Intercept) | 0.31 | 0.23 – 0.43 | **<0.001** |
| Condition [ACT] | 1.4 | 0.91 – 2.16 | 0.128 |
| Condition [CVS] | 1.99 | 1.31 – 3.02 | **0.001** |
| AP Position 1 | 0.91 | 0.60 – 1.37 | 0.641 |
| AP Position 2 | 0.91 | 0.59 – 1.41 | 0.674 |
| Sex [M] | 1.55 | 1.02 – 2.36 | **0.04** |
| Marker 1 | 0.64 | 0.58 – 0.71 | **<0.001** |
| Marker 2 | 1.25 | 1.13 – 1.37 | **<0.001** |
| Condition [ACT] × AP Position 1 | 1.18 | 0.65 – 2.14 | 0.585 |
| Condition [CVS] × AP Position 1 | 0.91 | 0.51 – 1.60 | 0.739 |
| Condition [ACT] × AP Position 2 | 1.02 | 0.56 – 1.89 | 0.938 |
| Condition [CVS] × AP Position 2 | 1.36 | 0.74 – 2.47 | 0.317 |
| Condition [ACT] × Sex [M] | 1.29 | 0.71 – 2.34 | 0.403 |
| Condition [CVS] × Sex [M] | 0.39 | 0.22 – 0.72 | **0.003** |
| AP Position 1 × Sex [M] | 1.06 | 0.59 – 1.91 | 0.836 |
| AP Position 2 × Sex [M] | 1.09 | 0.60 – 1.95 | 0.783 |
| Condition [ACT] × AP Position 1 × Sex [M] | 0.9 | 0.39 – 2.09 | 0.811 |
| Condition [CVS] × AP Position 1 × Sex [M] | 0.75 | 0.33 – 1.72 | 0.494 |
| Condition [ACT] × AP Position 1 × Sex [M] | 1.04 | 0.46 – 2.39 | 0.92 |
| Condition [CVS] × AP Position 2 × Sex [M] | 0.93 | 0.39 – 2.21 | 0.874 |

Observations : 224
Marginal R<sup>2</sup> / Conditional R<sup>2</sup> : 0.343 / 0.624

**Supplementary Table 4.** Linear-log model statistics for GFP expressing neurons.

| **Ratios** |  |  |  |
| --- | --- | --- | --- |
| *Predictors* | *Estimates* | *CI* | *p* |
| (Intercept) | 0.18 | 0.13 – 0.25 | **<0.001** |
| Condition [ACT] | 1.7 | 1.09 – 2.64 | **0.02** |
| Condition [CVS] | 2.54 | 1.65 – 3.90 | **<0.001** |
| AP Position 1 | 0.98 | 0.71 – 1.36 | 0.909 |
| AP Position 2 | 0.93 | 0.66 – 1.32 | 0.691 |
| Sex [M] | 1.61 | 1.05 – 2.47 | **0.031** |
| Condition [ACT] × AP Position 1 | 1.23 | 0.77 – 1.98 | 0.38 |
| Condition [CVS] × AP Position 1 | 0.97 | 0.62 – 1.51 | 0.887 |
| Condition [ACT] × AP Position 2 | 0.82 | 0.50 – 1.33 | 0.415 |
| Condition [CVS] × AP Position 2 | 1.07 | 0.66 – 1.73 | 0.776 |
| Condition [ACT] × Sex [M] | 1.37 | 0.74 – 2.51 | 0.309 |
| Condition [CVS] × Sex [M] | 0.26 | 0.14 – 0.49 | **<0.001** |
| AP Position 1 × Sex [M] | 1.01 | 0.64 – 1.61 | 0.954 |
| AP Position 2 × Sex [M] | 1.16 | 0.73 – 1.85 | 0.524 |
| Condition [ACT] × AP Position 1 × Sex [M] | 0.79 | 0.40 – 1.56 | 0.494 |
| Condition [CVS] × AP Position 1 × Sex [M] | 0.69 | 0.36 – 1.32 | 0.256 |
| Condition [ACT] × AP Position 1 × Sex [M] | 1.09 | 0.56 – 2.10 | 0.796 |
| Condition [CVS] × AP Position 2 × Sex [M] | 1.11 | 0.56 – 2.20 | 0.754 |

Observations : 76
Marginal R<sup>2</sup> / Conditional R<sup>2</sup> : 0.524 / 0.628

**Supplementary Table 5.** Linear-log model statistics for mCherry expressing neurons.

| *Predictors* | **Ratios** |  |  |
| --- | --- | --- | --- |
|  | *Estimates* | *CI* | *p* |
| (Intercept) | 0.32 | 0.19 – 0.53 | <0.001 |
| Condition [ACT] | 1.95 | 0.94 – 4.01 | 0.07 |
| Condition [CVS] | 2.61 | 1.28 – 5.33 | 0.009 |
| AP Position 1 | 0.93 | 0.67 – 1.30 | 0.674 |
| AP Position 2 | 1.02 | 0.71 – 1.46 | 0.933 |
| Sex [M] | 2.08 | 1.02 – 4.24 | 0.045 |
| Condition [ACT] × AP Position 1 | 1.31 | 0.80 – 2.15 | 0.275 |
| Condition [CVS] × AP Position 1 | 0.82 | 0.52 – 1.30 | 0.397 |
| Condition [ACT] × AP Position 2 | 0.94 | 0.57 – 1.56 | 0.815 |
| Condition [CVS] × AP Position 2 | 1.51 | 0.92 – 2.48 | 0.1 |
| Condition [ACT] × Sex [M] | 0.77 | 0.28 – 2.09 | 0.605 |
| Condition [CVS] × Sex [M] | 0.3 | 0.11 – 0.86 | 0.025 |
| AP Position 1 × Sex [M] | 0.88 | 0.55 – 1.41 | 0.585 |
| AP Position 2 × Sex [M] | 0.88 | 0.55 – 1.43 | 0.605 |
| Condition [ACT] × AP Position 1 × Sex [M] | 1.11 | 0.55 – 2.25 | 0.773 |
| Condition [CVS] × AP Position 1 × Sex [M] | 0.79 | 0.41 – 1.53 | 0.473 |
| Condition [ACT] × AP Position 1 × Sex [M] | 1.13 | 0.58 – 2.23 | 0.708 |
| Condition [CVS] × AP Position 2 × Sex [M] | 1.12 | 0.56 – 2.27 | 0.738 |

Observations : 76
Marginal R<sup>2</sup> / Conditional R<sup>2</sup>: 0.317 / 0.720

**Supplementary Table 6.** Linear-log model statistics for GFP/mCherry expressing neurons.

| *Predictors* | **Ratios** |  |  |
| --- | --- | --- | --- |
|  | *Estimates* | *CI* | *p* |
| (Intercept) | 0.59 | 0.34 – 1.01 | 0.054 |
| Condition [ACT] | 0.89 | 0.41 – 1.92 | 0.757 |
| Condition [CVS] | 1.2 | 0.57 – 2.51 | 0.627 |
| AP Position 1 | 0.74 | 0.43 – 1.27 | 0.269 |
| AP Position 2 | 0.8 | 0.44 – 1.43 | 0.439 |
| Sex [M] | 0.93 | 0.44 – 1.96 | 0.838 |
| Condition [ACT] × AP Position 1 | 0.97 | 0.43 – 2.19 | 0.931 |
| Condition [CVS] × AP Position 1 | 0.93 | 0.44 – 1.97 | 0.84 |
| Condition [ACT] × AP Position 2 | 1.49 | 0.65 – 3.42 | 0.343 |
| Condition [CVS] × AP Position 2 | 1.94 | 0.87 – 4.36 | 0.106 |
| Condition [ACT] × Sex [M] | 1.97 | 0.68 – 5.69 | 0.204 |
| Condition [CVS] × Sex [M] | 0.86 | 0.29 – 2.57 | 0.781 |
| AP Position 1 × Sex [M] | 1.36 | 0.62 – 3.00 | 0.433 |
| AP Position 2 × Sex [M] | 1.37 | 0.62 – 3.03 | 0.428 |
| Condition [ACT] × AP Position 1 × Sex [M] | 0.99 | 0.31 – 3.13 | 0.981 |
| Condition [CVS] × AP Position 1 × Sex [M] | 0.84 | 0.27 – 2.60 | 0.754 |
| Condition [ACT] × AP Position 1 × Sex [M] | 0.8 | 0.26 – 2.44 | 0.685 |
| Condition [CVS] × AP Position 2 × Sex [M] | 0.49 | 0.15 – 1.62 | 0.236 |

Observations : 72
Marginal R<sup>2</sup> / Conditional R<sup>2</sup>: 0.157 / 0.356

**Supplementary Figure 1.**
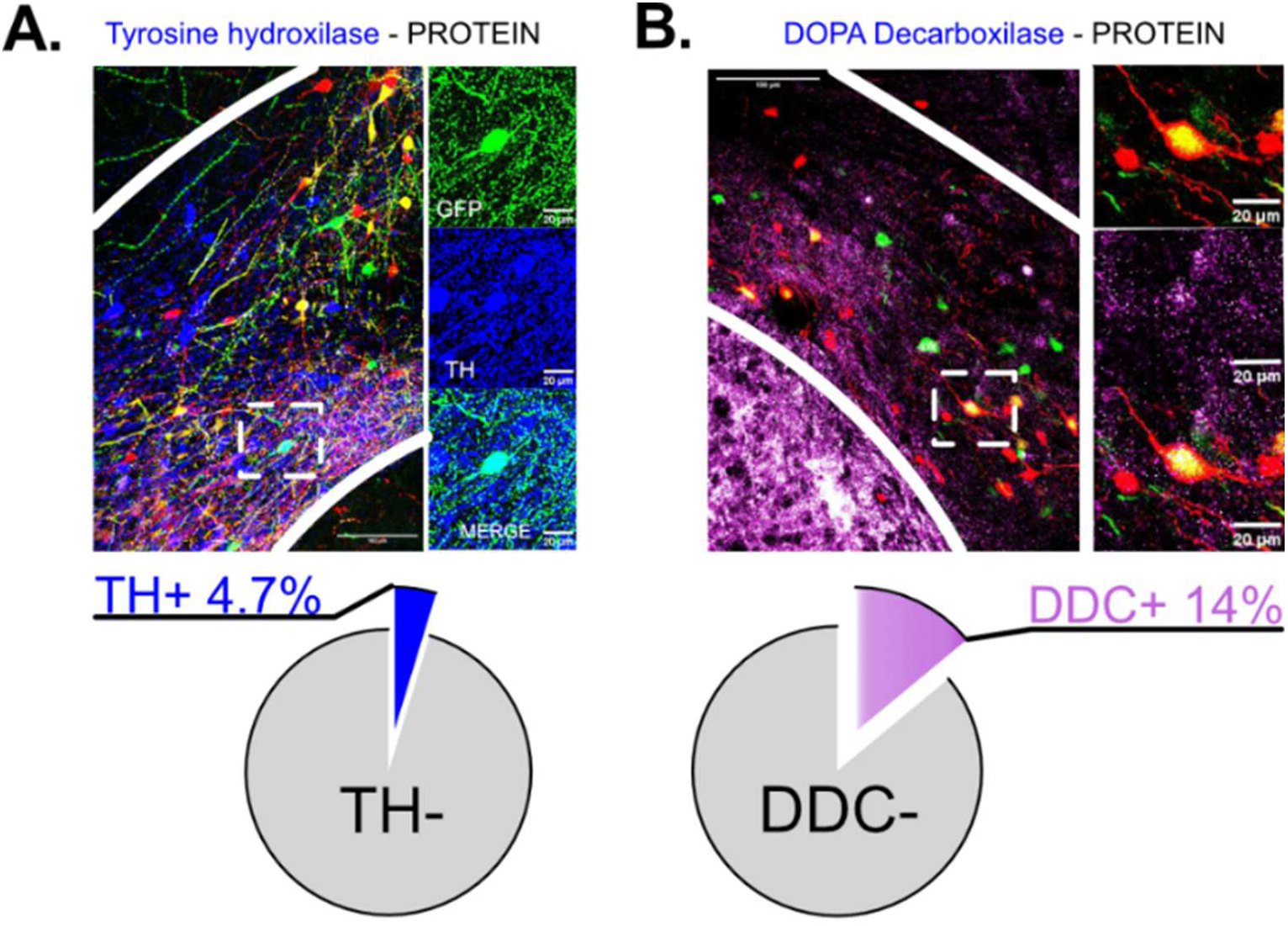
Immunohistochemistry-based quantification underestimates the identification of dopaminergic AAV-tagged cells. **A** Representative images of tyrosine hydroxylase (TH) immunohistochemistry in AAV-labeled cells, with insets showing the different fluorescent channels (TH in blue). Bottom: pie chart showing the proportion of TH+ and TH− cells. **B** Representative images of dopa decarboxylase (DDC) immunohistochemistry in AAV-labeled cells (DDC in purple), presented as in panel A.

**Supplementary Figure 2.**
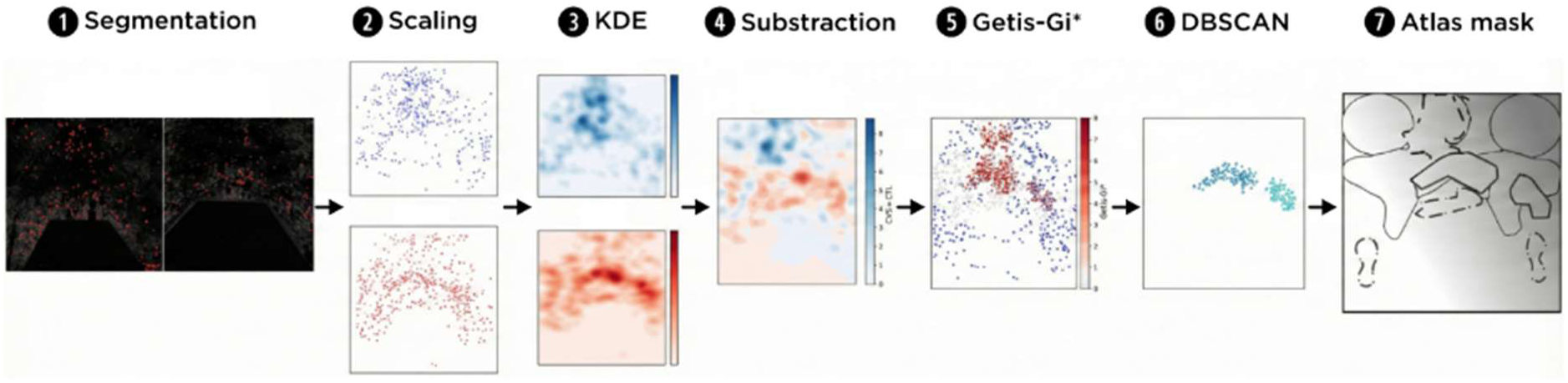
Computational pipeline for spatial hotspot analysis. Schematic representation of the computational workflow used to process c-Fos activity maps following acute and chronic stress. **1** Segmentation of c-Fos-positive nuclei from raw confocal images. **2** Normalization of spatial coordinates into a common [0,1] reference space. **3** Kernel density estimation (KDE) to generate continuous density maps. **4** Subtraction of control density maps from stress conditions to isolate condition-specific activity patterns. **5** Getis-Ord Gi* local spatial statistics to identify significant hotspots and coldspots. **6.** DBSCAN clustering to define the spatial boundaries of significant cell clusters. **7** Registration onto a standardized atlas mask for anatomical localization.

**Supplementary Figure 3.**
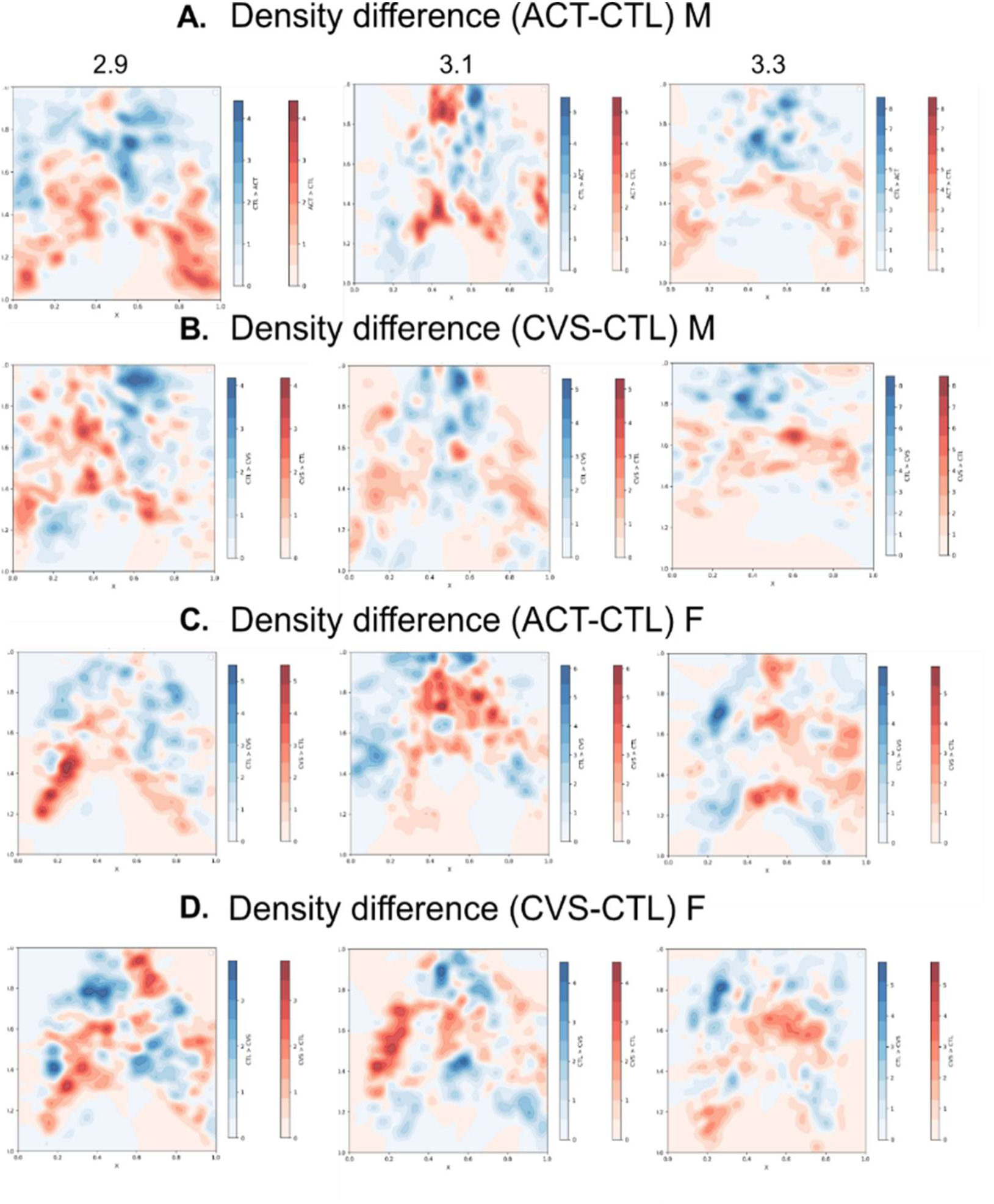
Spatial differences in c-Fos+ cell density across anterior–posterior levels of the ventral tegmental area in male and female mice. Heatmaps show relative differences in c-Fos^+^ cell density between **A** control (CTRL) and acute (ACT) stress conditions in males, **B** CTRL and chronic variable stress (CVS) conditions in males **C** CTRL and ACT stress conditions in females, and **D** CTRL and CVS conditions in females across AP levels of the VTA. Warmer colors (red) indicate regions with higher c-Fos^+^ density in stressed animals, whereas cooler colors (blue) indicate regions with higher density in control animals.

**Supplementary Figure 4.**
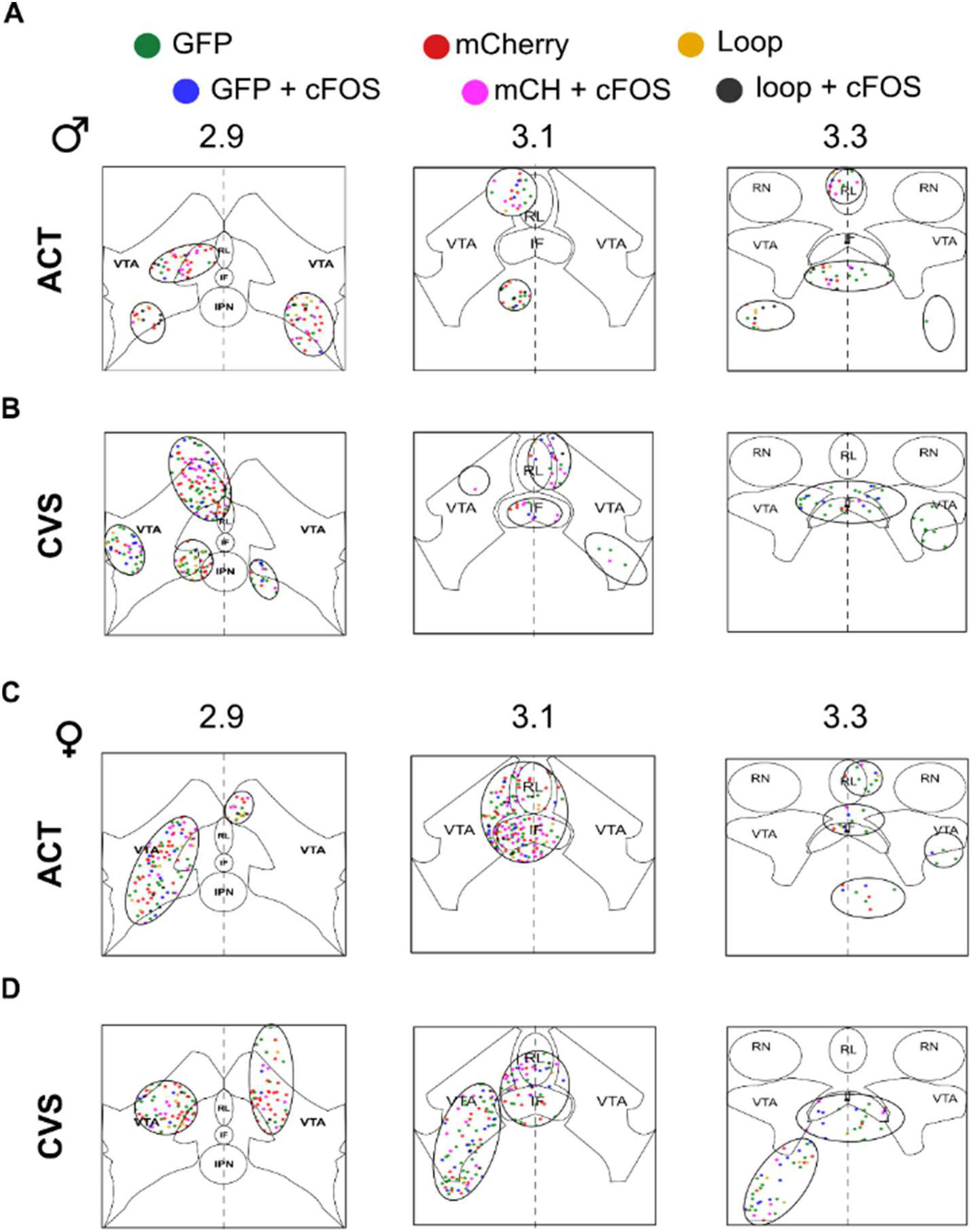
Spatial distribution and cellular composition of neuronal hotspots within the VTA. Representative coronal maps showing the cellular composition of hotspots identified through c-Fos density analysis across the AP axis (-2.9, -3.1, and -3.3 mm from Bregma). Data are shown for **A** acute stress in males, **B** chronic variable stress in males, **C** acute stress in females, and **D** chronic variable stress in females. Ellipses indicate identified hotspots within the ventral tegmental area (VTA), interfascicular nucleus (IF), and Rostral Linear Nucleus (RL). Colored markers indicate fluorescent cell identity: GFP^+^ (green), mCherry^+^ (red), and GFP^+^/mCherry^+^ cells (gold). c-Fos-positive cells within each population are shown in blue (GFP^+^/c-Fos^+^), pink (mCherry^+^/c-Fos^+^), and black (GFP^+^/mCherry^+^/c-Fos^+^). Additional anatomical landmarks include the interpeduncular nucleus (IPN) and red nucleus (RN).

